## Supplementary Tables and Figures revised for "Epistatic interactions between *PHOTOPERIOD-1, CONSTANS 1* and *CONSTANS 2* modulate the photoperiodic response in wheat"

**Supplementary Figure S1.** Shoot apical meristem (SAM) and spike development. Kronos-PI (*Ppd-A1a*), Kronos-PS (*Ppd-A1b*) and Kronos-*ppd1* loss-of-function mutant plants grown under LD (16 h light / 8 h darkness, top) and SD (8 h light / 16 h darkness, bottom). Bar is 200 μm in all figures. Samples are aligned by developmental stage (leaf number), but chronological time of dissections differed between LD and SD. Main tillers were dissected from three plants per genotype/time point and SAMs were photographed, but only one representative SAM of the three is included in the figure.

| Development | 4th leaf | 6th leaf | 8th leaf | 10th leaf | 12th leaf | 14th leaf |
|---|---|---|---|---|---|---|
| Time LD | 5 w | 6 w | 8 w | 10 w | 12 w | 14 w |
| Time SD | 5 w | 7 w | 9 w | 10.5 w | 13 w | 16 w |

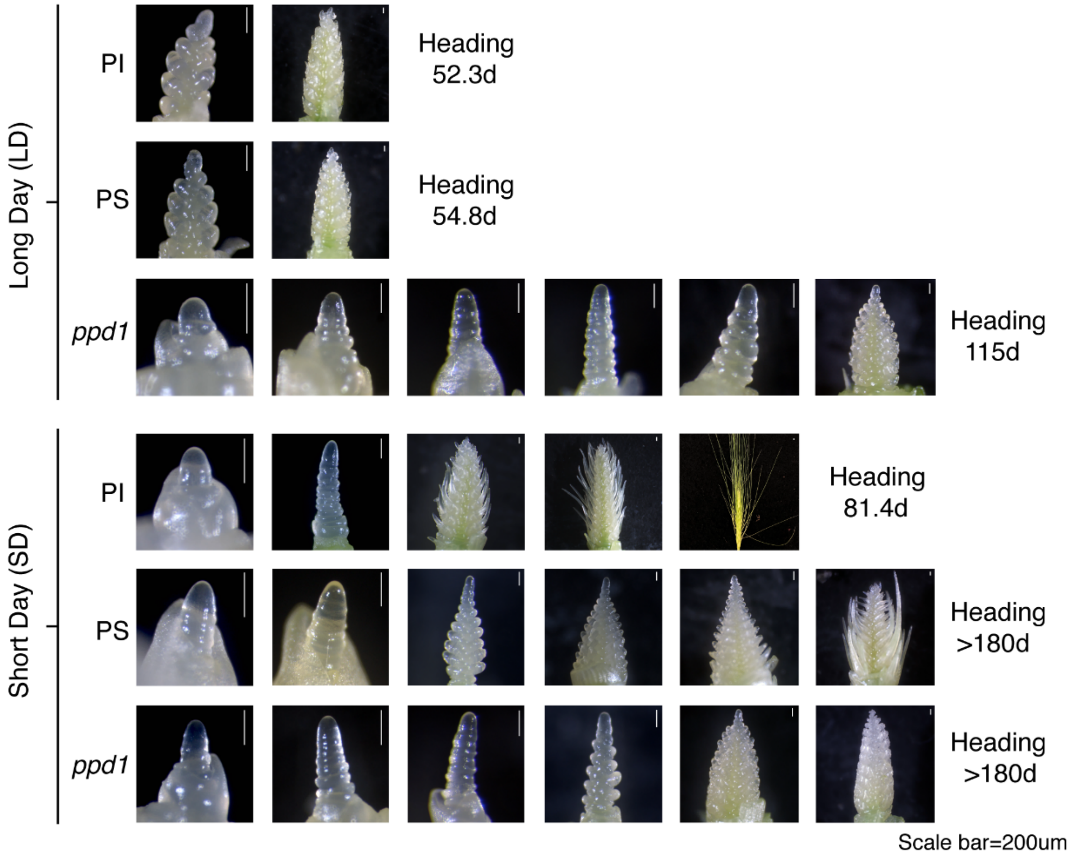

**Supplementary Figure S2**. Dissection of developing spikes. (**A**) Kronos-*ppd1* null mutant and (**B**) Kronos-PS control plants grown under SD and dissected 140 days (20 weeks) after sowing. Note the faster development of Kronos-PS relative to Kronos-*ppd1*-null. In both genotypes spikes failed to emerge before 180 days when the experiment was terminated.

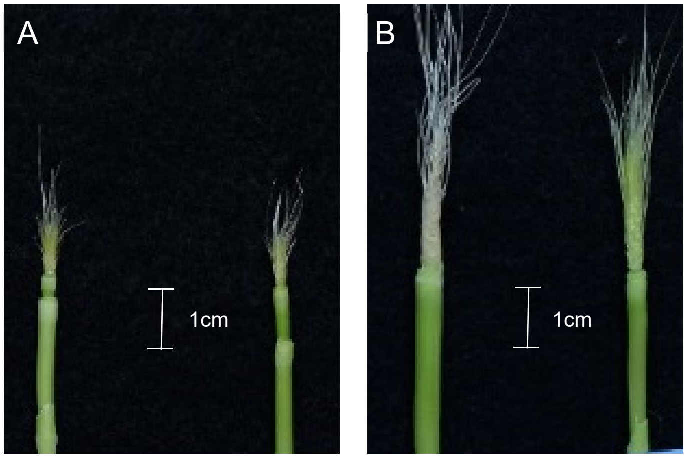


Kronos-*ppd1*-null Kronos-PS

**Supplementary Figure S3.** Effect of *ppd1*, *co1* and *co2* loss-of-function mutations and photoperiod on *CO1* and *CO2* transcript levels.  RNA samples were collected at ZT4 from leaves of six-week-old Kronos-PS and *ppd1* plants (with and without *co1* and *co2*). (**A**) *CO1* transcript levels. (**B**) *CO2* transcript levels. Dunnett's tests were used to compare the two mutants with the wild type. Transcript levels are expressed relative to *ACTIN* using the ΔCt method. Averages and standard errors of the means were calculated from a minimum of five biological replicates per genotype.

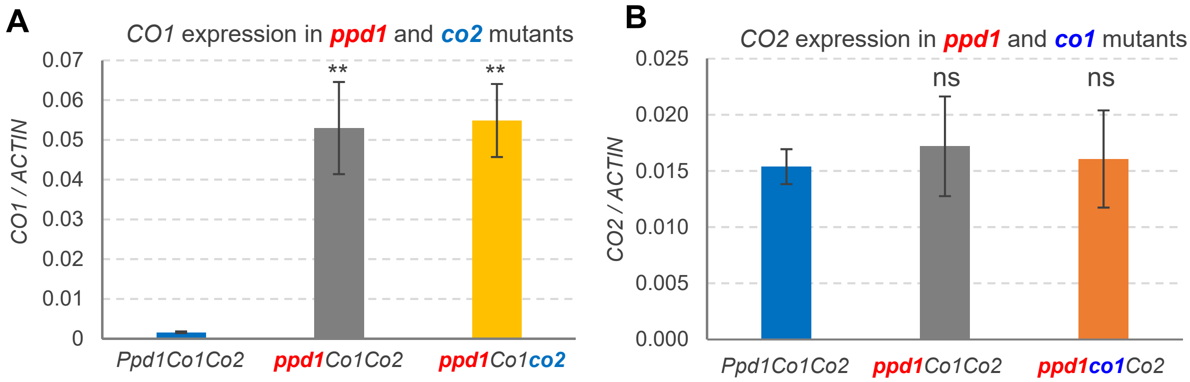

**Supplementary Figure S4.** Effect of *phyB*-null and *phyC*-null mutations in Kronos-PI on the transcriptional profiles of *CO1* and *CO2* under SD and LD. Results extracted from a published RNAseq study [1]. Samples were collected from the newest expanded leaves at ZT4 from 4w-old plants under LD (n = 4) and 8w-old plants under SD (n = 8) to synchronize wild type genotypes grown under different photoperiods to a similar early reproductive stages (W3 early spike development without terminal spikelet). Comparisons between LD and SD should be interpreted with caution, since photoperiod effects and chronological time effects are conflated. Factorial ANOVAS were performed separately for each gene and *P* values for photoperiod (SD vs LD), genotype (WT, *phyB*, *phyC*) and their interactions are indicated below the gene names.

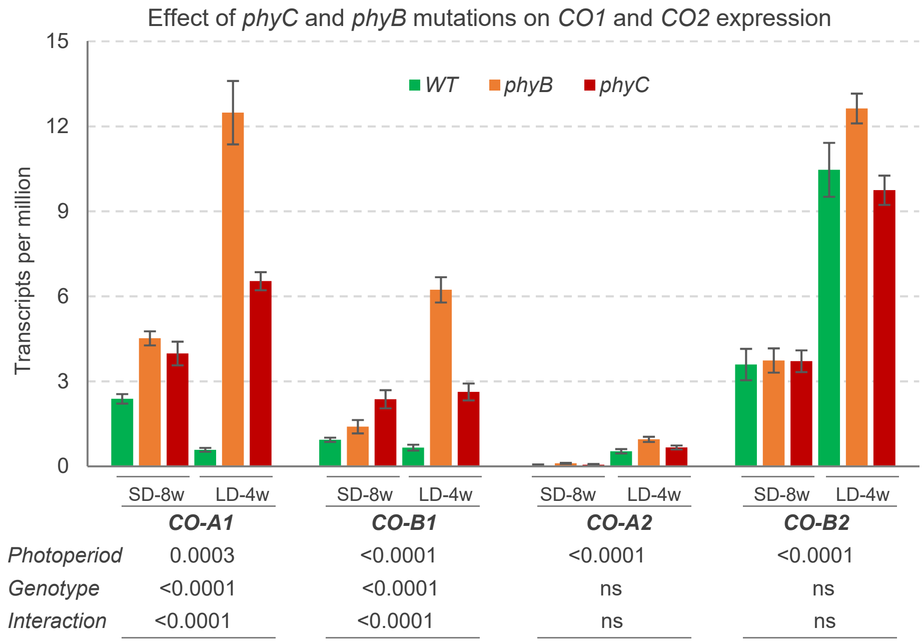

**Supplementary Figure S5.** Yeast-two-hybrid (Y2H) assays. Primers for cloning *PPD1* (from Kronos) and PHYB truncations (from *T. monococcum*) are listed in Supplementary Table S5. Primers and vectors for *CO1*, *CO2* and *VRN2* are described in [2] and those for *PHYC* and full-length *PHYB* in [3]. Transformants were selected on SD medium lacking leucine (L) and tryptophan (W) plates and re-plated on SD medium lacking L, W, histidine (H) and adenine (A) to test the interactions. Due to auto-activation, CO1 and CO2 can only be used as preys. Only N-PHYB can be used as bait without autoactivation, so this is the only PHYB clone tested for interactions with CO1 and CO2. For VRN2, we used the functional ZCCT1 paralog from *T. monococcum*. The PPD1-bait construct used in assays presented in this figure is the same as in Figure 7A showing a positive interaction with both CO1 and CO2.

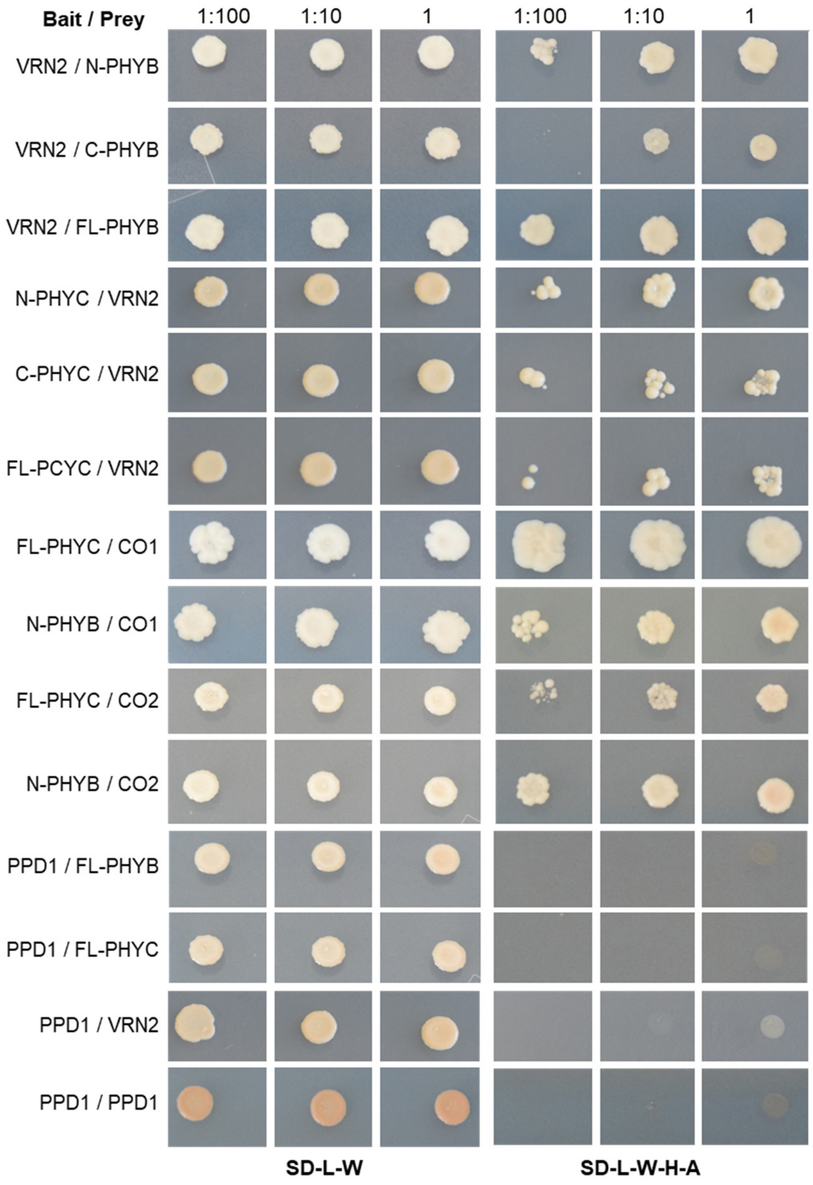

### Supplementary Tables

**Supplementary Table S1.** Genome-specific primer sequences and PCR conditions for TILLING. We sequenced genome-specific PCR products the first time to confirm amplification of the correct target.

| Gene name | Target | Primer name | Primer sequence (5' to 3') | Product (bp) | Ann. Temp (ºC) | Extension time | Enzyme |
|---|---|---|---|---|---|---|---|
| *CO1A* | Exon1 | CO1A-5P-F1 | ACATAGGCAGTGCATGAACACAT | 1174 | 59* | 1 m 30 s | *Hpy*188I |
| | | CO1A-5P-R2 | AGAAGTAGAAAAAGTTGAAGAAAGAG | | | | |
| *CO1A* | Exon 2 | CO1A-3P-F2 | CAATTCATCTCTAGGAAAGTAC | 955 | 54* | 1 m 30 s | - |
| | | CO1A-3P-R1 | CGTGCTATCTGAAACTATAAAC | | | | |
| *CO1B* | Exon 1 | CO1-5P-CF2 | CCACTGACACCCTACTATTAG | 1375 | 55* | 1 m 30 s | - |
| | | CO1B-5P-R2 | AGAAGTGGAAAAAGTTGAAGAAAGAA | | | | |
| *CO1B* | Exon 2 | CO1B-3P-F2 | CAATTCATCTCTAGGAAAGTAA | 987 | 54* | 1 m 30 s | *EcoR*V |
| | | CO1B-3P-R1 | CGTGCTATCTGAAACTATAAAT | | | | |
| *CO2A* | Exon 1 | CO2A-BBOX-F1 | TTCCAACACTGACTGCTTCTG | 1081 | 53* | 1 m 30 s | - |
| | | CO2-BBOX-CR1 | CTTGTATCCTAAGTGAGATGTGAC | | | | |
| *CO2A* | Exon 2 | CO2A-ZCCT-F1 | GGTTACAACTCTGGATGGTAG | 922 | 55 | 1 m 30 s | *EcoR*V |
| | | CO2-ZCCT-CR4 | GGACTATGTGGTTCACAATATG | | | | |
| *CO2B* | Exon 1 | CO2B-BBOX-F1 | TTCCAACACTGACTGCTCCCA | 1021 | 54* | 1 m 30 s | - |
| | | CO2-BBOX-CR1 | CTTGTATCCTAAGTGAGATGTGAC | | | | |
| *CO2B* | Exon 2 | CO2B-ZCCT-F1 | CTGTCCAACAGAATGTTTGAC | 931 | 54* | 1 m 30 s | - |
| | | CO2-ZCCT-CR3 | CTAACAGTAGAAGTCCCAACC | | | | |

*Touch-down protocol for PCR includes an additional 94 ºC for 5 m, 12 cycles of initial touch-down, with a reduction of 0.5 ºC per cycle (6 ºC total from final annealing T), then followed by 40 cycles of 94 ºC for 5 m, annealing temperature for 30 s and a final extension time of 7 m at 72 ºC.

**Supplementary Table S2.** Number of mutations detected in the targeted regions of wheat *CO1* and *CO2* homologs in the Kronos TILLING population.

| **Gene** | **Mutations (Total)** | **Non-synonymous** | **Synonymous** | **Splice or stop mutations** |
|---|---|---|---|---|
| *CO-A1* | 37 | 14 | 22 | 1 |
| *CO-B1* | 49 | 28 | 20 | 1 |
| *CO-A2* | 23 | 9 | 13 | 1 |
| *CO-B2* | 44 | 23 | 19 | 2 |
| Total | 153 | 74 | 74 | 5 |

**Supplementary Table S3**. Analysis of variance for heading time under long days (LD, 16 h light / 8 h darkness). This factorial ANOVA combined all four classes of *CO1* and *CO2* wild type and mutant alleles in photoperiod sensitive (PS, *Ppd-A1b*, three experiments) and photoperiod insensitive backgrounds (PI, *Ppd-A1a*, two experiments). For the statistical analyses, we used experiments as blocks nested within *PPD1* classes.

Dependent Variable: Heading time

| Source | DF | Sum of Squares | Mean Square | *F* Value | *P* > F |
|---|---|---|---|---|---|
| Model | 10 | 948.953 | 94.895 | 38.04 | <.0001 |
| Error | 51 | 376.658 | 2.494 | | |
| Corrected Total | 61 | 1325.611 | | | |

$R^2$=0.716

**S3.A**. Overall 3-way ANOVA.

| Source | DF | Type III SS | Mean Square | *F* Value | *P* > F |
|---|---|---|---|---|---|
| *Experiment* | 3 | 55.847 | 18.616 | 7.46 | 0.0001 |
| *PPD1* | 1 | 408.755 | 408.755 | 163.87 | <.0001 |
| *CO1* | 1 | 173.578 | 173.578 | 69.59 | <.0001 |
| *CO2* | 1 | 33.081 | 33.081 | 13.26 | 0.0004 |
| *CO1*CO2* | 1 | 2.250 | 2.250 | 0.90 | 0.3437 |
| *PPD1*CO1* | 1 | 55.066 | 55.066 | 22.08 | <.0001 |
| *PPD1*CO2* | 1 | 78.353 | 78.353 | 31.41 | <.0001 |
| *PPD1*CO1*CO2* | 1 | 20.670 | 20.670 | 8.29 | 0.0046 |

**S3.B**. 2-way ANOVA *CO1* x *CO2* in PS (Main text, Fig. 2B).

| Source | DF | Type III SS | Mean Square | *F* Value | *P* > F |
|---|---|---|---|---|---|
| Exp | 2 | 55.073 | 27.536 | 10.55 | <.0001 |
| CO1 | 1 | 249.762 | 249.762 | 95.67 | <.0001 |
| CO2 | 1 | 5.653 | 5.653 | 2.17 | 0.1448 |
| CO1*CO2 | 1 | 21.494 | 21.494 | 8.23 | 0.0052 |

**S3.C**. 2-way ANOVA *CO1* x *CO2* in PI (Main text, Fig. 2C).

| Source | DF | Type III SS | Mean Square | *F* Value | *P* > F |
|---|---|---|---|---|---|
| Exp | 1 | 0.775 | 0.775 | 0.33 | 0.5668 |
| CO1 | 1 | 14.386 | 14.386 | 6.16 | 0.0157 |
| CO2 | 1 | 92.724 | 92.724 | 39.69 | <.0001 |
| CO1*CO2 | 1 | 4.036 | 4.036 | 1.73 | 0.1934 |

**Supplementary Table S4**. Analysis of variance for heading time under short days (SD, 8 h light / 16 h darkness). All four allelic combinations for *CO1* and *CO2* wild type and mutant alleles were analyzed in a photoperiod insensitive background (Kronos-PI).

Dependent Variable: Heading time

| Source | DF | Type III SS | Mean Square | *F* Value | *P* > F |
|---|---|---|---|---|---|
| Model | 3 | 1099.363 | 366.454 | 28.61 | <.0001 |
| Error | 35 | 448.226 | 12.806 | | |
| Corrected Total | 38 | 1547.590 | | | |

$R^2$ 0.710

| Source | DF | Type III SS | Mean Square | *F* Value | *P* > F |
|---|---|---|---|---|---|
| Main effects | | | | | |
| *CO1* | 1 | 324.801 | 324.801 | 25.36 | <.0001 |
| *CO2* | 1 | 574.507 | 574.507 | 44.86 | <.0001 |
| *CO1 x CO2* | 1 | 131.177 | 131.177 | 10.24 | 0.0029 |
| Simple effects | | | | | |
| *CO1-co1* in *CO2* WT | 1 | 20.056 | 20.056 | 1.57 | 0.2191 |
| *CO1-co1* in *co2* mut | 1 | 470.028 | 470.028 | 36.70 | <.0001* |
| *CO2-co2* in *CO1* WT | 1 | 76.634 | 76.634 | 5.98 | 0.0196 |
| *CO2-co2* in *co1* mut | 1 | 641.479 | 641.479 | 50.09 | <.0001* |

*CO1 and CO2 are highly significant only in the presence of the other mutant allele

**Supplementary Table S5**. Primers used in the qRT-PCR and Y2H experiments.

| Primer name | Primer Sequence | Genome | Efficiency | Reference |
|---|---|---|---|---|
| **qRT-PCR** | | | | |
| SYBR-ACTIN-F | ACCTTCAGTTGCCCAGCAAT | | 98 % | [4] |
| SYBR-ACTIN-R | CAGAGTCGAGCACAATACCAGTTG | | | |
| TaFT1_AB_qPCR_F | CAGCAGCCCAGGGTTGAG | A & B | 97 % | [5] |
| TaFT1_AB_qPCR_R | ATCTGGGTCTACCATCACGAGTG | | | |
| Vrn1-Ex5-6-F2 | AAGAAGGAGAGGTCACTGCAGG | A & B | >96 % | [5] |
| Vrn1-Ex8-R37 | GGCTGCACTGCCGCA | | | |
| Vrn2_ZCCT2_F | CCACCATCGTGCCATTCT | A & B | >96 % | [4] |
| Vrn2_ZCCT2_R | CCCACCATCATCTCTGTATCAA | | | |
| PPD1_SYB-F3 | CGGCATTCACGAGGTACAATAC | A & B | 97.6% | [3] |
| PPD1_SYB-R3 | GAGCCTTGCTTCATCTGAGCG | | | |
| AC-CO1-AB-F3 | CACATCAGAGTGGTTATGC | A & B | 94.0 % | [3] |
| AC-CO1-AB-R3 | GGACTGGACCGTATTGTC | | | |
| AC-CO2-AB-SYB-F4 | AAGGGTGTGAGTGTGTAG | A & B | 96.0 % | [3] |
| AC-CO2-AB-SYB-R5 | GATATGTCATTGCTGATGGAAG | | | |
| **Y2H** [1] | | | | |
| PPD1-F [2] | CATATGGACCGTCATCACCAGCAG | | | |
| PPD1-R | GAATTCTCTCTCCACGGCAGCCGGCGG | | | |
| N-TmPHYB-Y2H-F [3] | **CCGAATTC**ATGGCCTCGGGAAGCCGCGC | | | |
| N-TmPHYB-Y2H-F | **TCCCCCCGGG**TGCATCTCTGAAGGAGTCCCG | | | |
| C-TmPHYB-Y2H-F | CCGAATTCGGAGAGGGCACTAGTAACTC | | | |
| C-TmPHYB-Y2H-F | CCATCGATGCTCCGATCCCTACTTTCTG | | | |

[1] Y2H primers for cloning *CO1*, *CO2* and *VRN2* have been described in [2] and those for *PHYC* full-length and truncations and *PHYB* ful-length have been described in [3].

[2] *PPD1* full-length coding region was cloned from Kronos.

[3] *PHYB* clones are from *T. monococcum.* Underlined bases indicate restriction sites used in cloning.
